# Morphological and Functional Effects of Cytoskeletal and Ion-Channel Agents on the Protoscolex of *Echinococcus granulosus sensu lato*

**DOI:** 10.64898/2026.04.06.716494

**Authors:** Mónica Patricia A. Carabajal, María José Fernández Salom, Luciano José Martínez, David Di Lullo, Enzo Rubén Marcial, Virginia Helena Albarracín, Horacio Fabio Cantiello

**Affiliations:** Laboratorio de Canales Iónicos, Instituto Multidisciplinario de Salud, Tecnología y Desarrollo (IMSaTeD, CONICET-UNSE)-Facultad de Agronomía y Agroindustria, Universidad Nacional de Santiago del Estero, Santiago del Estero, Argentina; Centro Integral de Microscopía Electrónica (CIME-UNT-CONICET), CCT CONICET NOA Sur, Universidad Nacional de Tucumán, Tucumán, Argentina; Cátedra de Biología Molecular e Ingeniería Genética, Instituto de Salud y Calidad de Vida, Universidad San Pablo-T, Lules, Tucumán, Argentina; Cátedra de Microscopía Electrónica (Métodos Auxiliares I), Especialización Anatomía Patológica, Facultad de Medicina, Universidad Nacional de Tucumán, Tucumán, Argentina; Cátedra de Biología Molecular, Facultad de Ciencias Naturales e Instituto Miguel Lillo, Universidad Nacional de Tucumán, Tucumán, Argentina

**Author notes:** **Correspondence**: Cantiello, HF, Instituto Multidisciplinario de Salud, Tecnología y Desarrollo (IMSaTeD, CONICET-UNSE)-Predio FAyA, RN 9, km 1125, Villa El Zanjón, Santiago Del Estero, Argentina, 4206.

**Keywords:** Echinococcus granulosus, Protoscoleces, Motility, Viability, Cytoskeleton, Ion channels, Ultrastructure

## Abstract

Helminthiases remain a major global health burden, and limitations of current anthelmintic therapies highlight the need for new pharmacological targets. In this study, we examined the effects of ion channel and cytoskeletal modulators on bovine lung protoscoleces (PSCs) of *Echinococcus granulosus* sensu lato. Compounds acting on ion channels (praziquantel, amiloride, and amlodipine) and cytoskeletal components (albendazole and cytochalasin D) were evaluated using a semi-automated motility assay, methylene blue exclusion to assess viability, and scanning electron microscopy (SEM) to characterize structural damage. All compounds produced concentration-dependent reductions in motility. Amlodipine was the most potent inhibitor of motility, whereas praziquantel and cytochalasin D produced pronounced tegumental alterations. Amiloride was the least potent compound at both endpoints and produced the mildest ultrastructural lesions. In all cases, movement was abolished at concentrations well below those required to reduce viability. Once their shared dependence on concentration was accounted for, the association between the two endpoints disappeared, indicating that paralysis and loss of viability are largely distinct responses. Motility did not recover in any compound after 24 h of drug washout. SEM analysis revealed extensive tegumental collapse, surface erosion and microtrichial loss in PSCs exposed to cytoskeletal and calcium-modulating agents. These findings highlight cytoskeletal organization and calcium-dependent ion fluxes as key physiological vulnerabilities in *E. granulosus*.

## 1. Introduction

Cystic echinococcosis (CE) is a neglected zoonotic disease of worldwide concern. It is caused by larval stages of species of the *Echinococcus granulosus* sensu lato (s.l.) complex, affecting humans and livestock on every continent studied to date, except Antarctica (Alvarez Rojas et al., 2014; Deplazes et al., 2017; WHO, 2023). Infection occurs following ingestion of parasite eggs from contaminated environments, leading to the development of hydatid cysts, primarily in the liver and lungs (Romig et al., 2017).

Within the hydatid cyst, protoscoleces (PSCs) emerge as the key infectious stage. They are enclosed by a laminated and germinal layer that provides structural support and facilitates immune evasion, thereby promoting parasite survival (Morseth, 1967). Cysts expand slowly and may remain asymptomatic for years, with clinical manifestations emerging as they enlarge and compress surrounding tissues (Kern et al., 2017). Surgical removal of the entire cyst is considered curative. However, accidental spillage of viable PSCs during surgery frequently results in secondary infection and disease recurrence (Prousalidis et al., 2012; Sielaff et al., 2001).

Pharmacological treatment of the disease relies mainly on benzimidazoles and, in some cases, praziquantel. Yet, both treatments present important limitations, including incomplete efficacy, variable clinical outcomes, adverse side effects, and the risk of emerging resistance (Doenhoff et al., 2008; Krämer et al., 2025). These shortcomings underscore the need to explore new therapeutic targets that exploit parasite-specific vulnerabilities. Among the most promising candidates are the cytoskeleton and ion channels, two systems essential for parasite physiology yet sufficiently divergent from their host counterparts to serve as selective drug targets (Perez-Serrano et al., 1995; Sasaki et al., 2002; Wolstenholme, 2011).

The parasite cytoskeleton plays a fundamental role in maintaining PSC morphology, motility, nutrient uptake, and tegumental integrity (Chen et al., 2008). Its divergence from mammalian cytoskeletal proteins has already been successfully exploited by benzimidazole carbamates, which selectively bind parasite β-tubulin (Guarnaschelli and Koziol, 2025). Beyond microtubules, actin filaments also support attachment and movement through specialized structures such as suckers and the rostellum, suggesting that disruption of actin dynamics could profoundly compromise parasite viability (La-Rocca et al., 2019).

Ion channels also govern essential physiological processes, including neuromuscular coordination, osmoregulation, and signal transduction of the parasite (Choudhary et al., 2022; Wolstenholme, 2011). Targeting ion transport has already proven highly effective in flatworm control. For instance, praziquantel’s mode of action implicates Ca²⁺-permeable channels as pivotal drug targets (Park and Marchant, 2020; Nogueira et al., 2022) thereby is linking calcium entry to rapid muscle contraction, tegumental disruption, and paralysis. Moreover, genomic analysis of *E. granulosus* has identified genes encoding Na⁺-like channel subunits, suggesting that other cation channels may play important functional roles, representing underexplored pharmacological opportunities (Zheng et al., 2013).

In this study, we evaluated the *in vitro* effects of cytoskeletal and ion-channel modulators on bovine lung PSCs of *E. granulosus*. Using motility assays, methylene blue viability staining, and scanning electron microscopy (SEM), we generated dose–response curves, calculated IC₅₀ values, and assessed structural damage at pharmacologically relevant doses. By integrating functional and morphological readouts, our work provides comparative insights into how disruption of cytoskeletal dynamics and cation-channel activity compromise PSC motility and viability, highlighting both systems as promising therapeutic targets for cestode infections.

## 2. Materials and methods

### 2.1. Chemicals

Dimethyl sulfoxide (DMSO), penicillin-streptomycin solution, cytochalasin D (CD), praziquantel (PZQ), albendazole (ABZ), amiloride (AMI), and amlodipine (AML) were obtained from Sigma-Aldrich Co. (St. Louis, MO, USA). Dulbecco’s Modified Eagle’s medium (DMEM) was supplied by Thermo Fisher Scientific (Waltham, MA, USA).

### 2.2. Parasites and incubation media

Protoscoleces of *Echinococcus granulosus* s.l. were extracted under sterile conditions from intact cysts of naturally infected cattle lungs. Briefly, bovine lungs were obtained from a local abattoir and aseptically collected PSCs were washed three times with sterile PBS (pH 7.2), as previously reported (Carabajal et al., 2022). After the last sedimentation, PSCs were resuspended in prewarmed culture medium (DMEM containing 0.4% (w/v) glucose, 100 IU/ml of penicillin, and 100 µg/ml of streptomycin) and incubated at 37°C in a humidified incubator with 5% CO_2_. Parasites were cultured for at least 24 h until the motility assay was performed.

### 2.3. Motility assessment

The assays were conducted in sterile 96-well flat-bottom microplates. Cultured parasites were dispensed into each well to obtain 50–60 PSCs per well. Test compounds were dissolved in DMSO, and 10 µL of each stock solution was added to reach final concentrations ranging from 10⁻¹⁰ M to 10⁻² M. Solvent and untreated controls were included in all experiments. Microplates were incubated under the same conditions described in Section 2.2.

PSCs motility was evaluated using a live-cell imaging system (Zeiss AxioScope A1 microscope equipped with an Axiocam 105 5.0 MP camera) and ZEISS ZEN Microscopy Software. For each well, two sequential images were captured at 10-second intervals under 40× magnification (Figure 1). Motility was quantified by calculating pixel-based displacement between the two frames in ImageJ v1.46, following the macro described by Ritler et al. (2017), with minor modifications. The motility index for each well was expressed as the percentage of movement relative to the DMSO control. To evaluate the reversibility of drug-induced motility inhibition, PSCs were first exposed to each compound for 24 h and then thoroughly washed with pre-warmed sterile PBS to remove the residual drug. Then, PSCs were further resuspended in fresh drug-free DMEM and incubated for an additional 24 h under identical culture conditions. Motility was quantified immediately after the initial 24 h exposure and again following the 24 h recovery period in the absence of drug.

**Figure 1.**
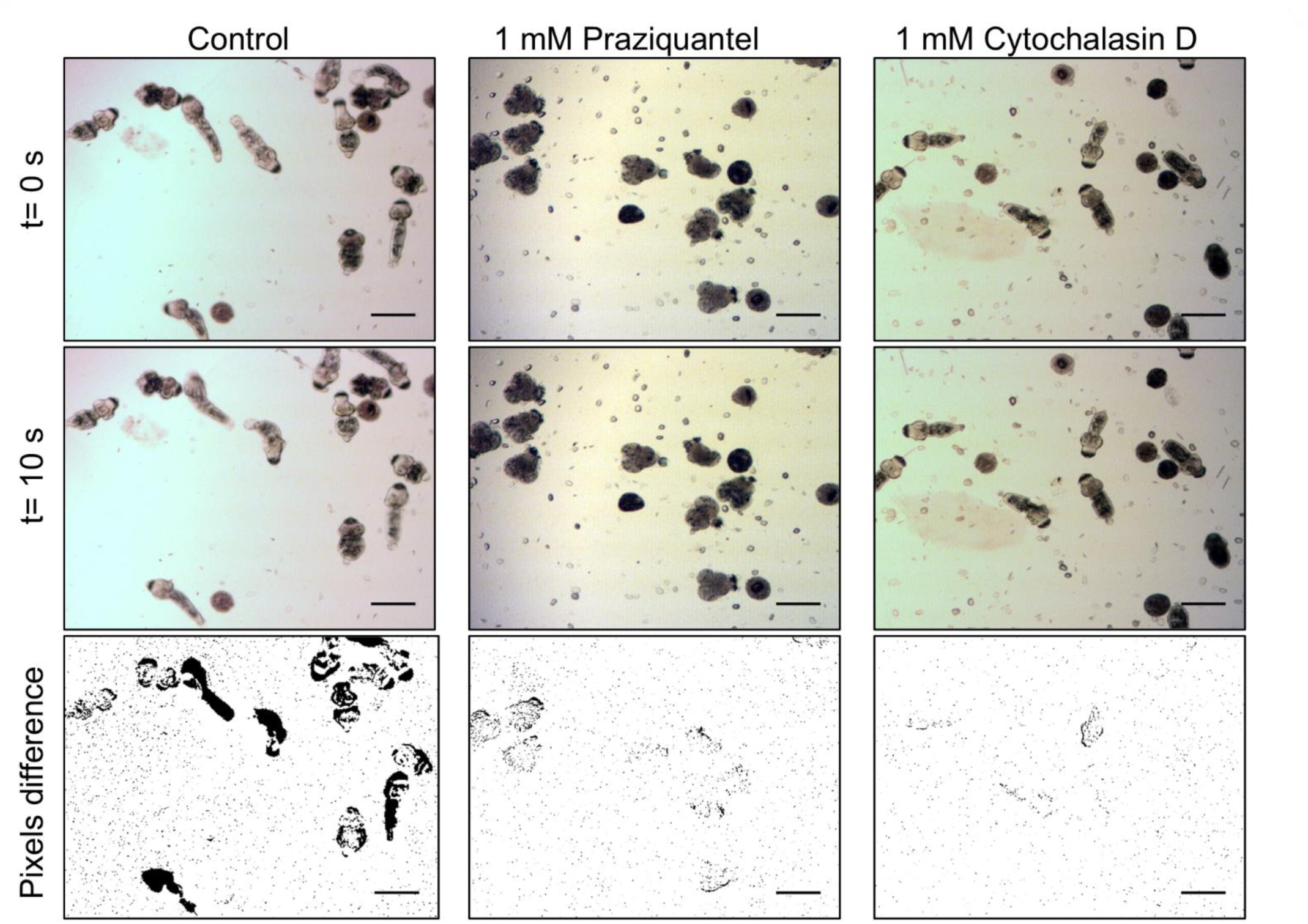
Representative images of motility analysis in *E. granulosus* PSCs. Photomicrographs illustrate the motility assessment under different experimental conditions. For each treatment, two consecutive frames separated by a 10-second interval are shown (first and second rows), allowing visualization of parasite displacement over time. Columns correspond to the experimental groups: untreated control (left), praziquantel-treated PSCs (middle), and cytochalasin D–treated PSCs (right). The bottom row shows the processed difference images obtained by pixel subtraction between the two frames, which highlight areas of movement and were used for quantitative motility analysis. Scale bar = 200 µm.

### 2.4. Viability assessment

Parasite viability was determined by the Methylene Blue (MB) exclusion assay, which remains one of the most commonly used methods because it reflects tegumental membrane integrity and provides a rapid, reproducible estimate of parasite survival (Casado et al., 1986; Ritler et al., 2017). Briefly, after 24 h of incubation with each tested compound, 0.1% MB solution was added to each well and mixed gently. After 5 min at room temperature, samples were examined under an inverted microscope (40×). Viable PSCs were identified by exclusion of the dye and preservation of normal morphology, whereas non-viable PSCs exhibited uniform blue staining due to loss of membrane integrity.

### 2.5. Scanning electron microscopy (SEM)

To evaluate tegumental morphological damage induced by drug treatment, SEM was performed as described in Rasuk et al. (2017), with slight modifications for parasite specimens. Briefly, PSCs were washed in PBS (pH 7.4) and fixed with Karnovsky’s fixative, containing 2.66% w/v paraformaldehyde and 1.66% w/v glutaraldehyde in 0.1 M phosphate buffer (pH 7.2). After fixation, PSCs were washed with 0.1 M phosphate buffer twice and dehydrated in a graded ethanol series (50%, 70%, 90%, and 100%), followed by 100% acetone. Critical-point drying was performed using a Denton Vacuum DCP-1. The samples were then mounted on aluminum stubs and subjected to gold sputtering (JEOL model JFC-1100). Control and treated PSC were observed under Zeiss Supra 55VP scanning electron microscope (Carl Zeiss NTS GmbH, Germany), at the Electron Microscopy Core Facility (CIME-CONICET-UNT).

### 2.6. Statistical analysis

Motility and viability values were expressed as a percentage of the control vehicle and fitted by nonlinear regression to a four-parameter logistic (4PL) model:

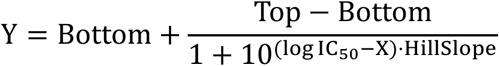

Because responses were normalized to the control, the upper asymptote (Top) was constrained to 100%. The Bottom was initially left free, bounded to [0, 100], and was retained as a free parameter when it was significantly greater than zero in a one-sided Wald test (p < 0.05); otherwise it was constrained to zero. Absolute IC_50_ and LC_50_ values were obtained by solving the fitted curve for a 50 % response rather than from the midpoint parameter, which coincide only when Bottom is zero. Motility dose-response curves were also fitted to a two-component model and compared with the 4PL by AICc,

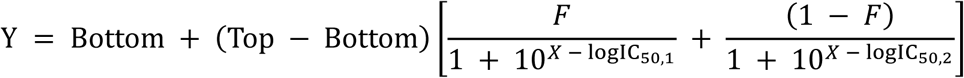

Correlation between motility and viability data was analyzed using Spearman’s rank (rₛ) correlation coefficient. The unit of analysis was the mean of each compound–concentration combination. The five per-compound tests were corrected by the Holm–Bonferroni procedure. To separate the association between endpoints from their shared concentration dependence, partial Spearman correlations were computed on ranks after regressing out log₁₀[concentration], and compound identity where indicated.

Biphasic behavior was tested by fitting a continuous segmented regression of viability (V) on motility inhibition (*M*),

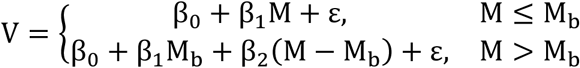

Statistical significance was set at p < 0.05 (Supplementary Table 1).

The change in motility was computed as Δ = M2 – M1, where M2 is motility 24 h after washout and M1 motility after 24 h of drug treatment; both measurements originate from the same well, thus pairing each post-washout value with its own pre-washout value. Treatment–concentration combinations for which mean post-washout viability fell below 15% were excluded from all analyses, because the motility readout becomes unreliable when almost no viable organisms remain. The dose dependence of recovery was modeled separately for each compound as Δ ∼ log10[dose] + experiment, with experiment included as a fixed effect. All analyses were performed in R 4.4.1 (R Core Team, 2024).

## 3. Results

### 3.1. Concentration-dependent effects on motility and viability

Motility and viability of *E. granulosus* s.l. PSCs were evaluated in response to increasing concentrations of ion-channel and cytoskeleton-modulating compounds. All five compounds inhibited PSCs motility in a concentration-dependent manner after 24 h of incubation, but the shape of the dose–response relationship differed markedly among them (Figure 2). PZQ and CD fell to an intermediate level of motility and remained there across several orders of magnitude before declining again at the highest concentrations, a profile that a single sigmoidal curve cannot follow; amlodipine showed a similar but less pronounced pattern, whereas amiloride and albendazole declined continuously. AML was the most potent compound, nearly abolishing movement at nanomolar concentrations (IC_50_ = 0.021 µM), and AMI the least potent (IC_50_ = 39.7 µM), with PZQ, ABZ and CD in between (IC_50_ = 0.03–0.43 µM; Table 1).

**Figure 2.**
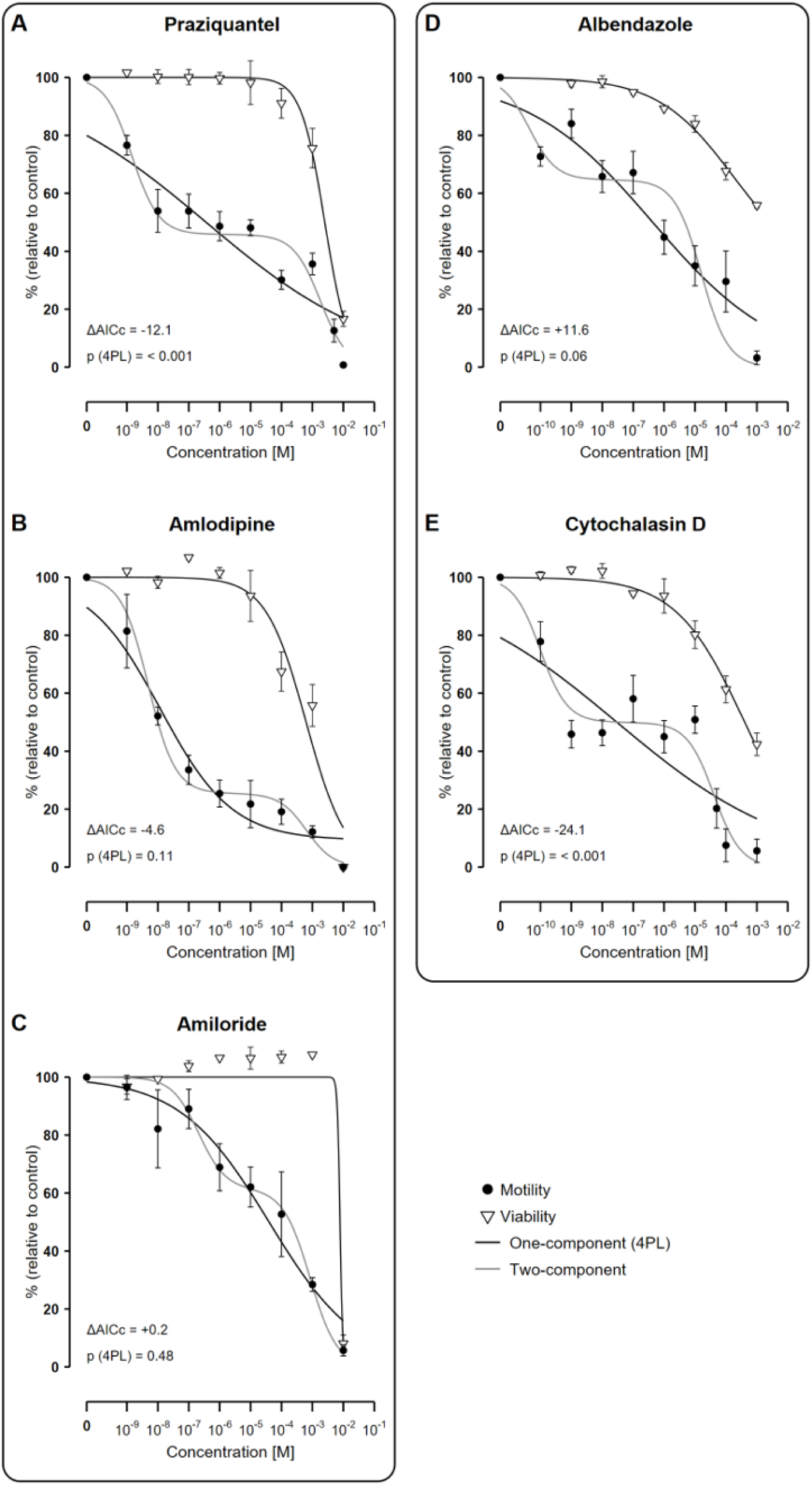
Concentration-dependent effects on motility and viability of *E. granulosus* PSCs. Dose–response curves of parasite motility (circles) and viability (triangles) after 24 h of exposure to (A) praziquantel, (B) amlodipine, (C) amiloride, (D) albendazole and (E) cytochalasin D. Data represent mean ± SEM from three independent experiments performed in duplicate; Black lines are four-parameter logistic fits, shown for both endpoints; those for motility are the fits reported in Table 1. Grey lines are two-component fits to the motility data, with both Hill slopes fixed at −1. Each panel gives ΔAICc, which compares the two motility models against each other and favours the two-component fit when negative, and the lack-of-fit p value of the four-parameter logistic, which compares it against experimental variability. Fitted parameters are given in Table 1 and Supplementary Table 1.

**Table 1.** Summary of dose–response parameters for viability and motility inhibition of *E. granulosus* protoscoleces after 24 h exposure to cytoskeletal disruptors and ion channel agents.

|  | Motility |  |  | Viability |  |  |
| --- | --- | --- | --- | --- | --- | --- |
| | IC<sub>50</sub>, μM [95% CI] | E<sub>max</sub> (%) | R<sup>2</sup> | LC<sub>50</sub>, mM [95% CI] | E<sub>max</sub> (%) | R<sup>2</sup> |
| **Cytoskeletal Modulators** |  |  |  |  |  |  |
| Albendazole | 0.428 [0.133–1.33] | 96.8 ± 2.4 | 0.74 | 2.87 [1.23–n.d.]<sup>a</sup> | 44.1 ± 1.0 | 0.96 |
| Cytochalasin D | 0.0295 [0.00369–0.157] | 94.4 ± 4.0 | 0.55 | 0.405 [0.257–0.679] | 57.7 ± 3.8 | 0.96 |
| **Ion Channel Agents** |  |  |  |  |  |  |
| Praziquantel | 0.364 [0.0826–1.31] | 99.2 ± 0.3 | 0.69 | 2.51 [1.91–3.30] | 83.4 ± 2.6 | 0.97 |
| Amlodipine | 0.0207 [0.00866–0.0549] | 100.0 ± 0.0 | 0.78 | 0.627 [0.332–1.16] | 100.0 ± 0.0 | 0.91 |
| Amiloride | 39.7 [13.8–114] | 94.3 ± 1.9 | 0.73 | 7.84 [3.25–8.44] | 91.9 ± 3.0 | 0.98 |

IC<sub>50</sub>, absolute half-maximal inhibitory concentration; LC<sub>50</sub>, absolute half-maximal lethal concentration. E<sub>max</sub> is the mean observed inhibition at the highest tested concentration ± SEM. Confidence intervals were obtained by profile likelihood.
<sup>a</sup> LC<sub>50</sub> is obtained by extrapolation, and its upper 95% limit is not constrained by the data; n.d., not determined.

This difference in curve shape was then examined formally. Hill slopes of the 4PL fits ranged from −0.15 to −0.37, with no 95 % confidence interval reaching −1, the value expected for a single-site saturable process, and the fits accounted for less of the variance in motility (R² = 0.55–0.78) than in viability (R² = 0.91–0.98); viability curves showed no comparable flattening (Supplementary Table 1). A lack-of-fit test against pure error indicated that the 4PL was inadequate only for PZQ (F₇,₃₆ = 5.40, p = 0.0003) and CD (F₇,₃₇ = 7.19, p = 1.9 × 10⁻⁵). For these two compounds, and for AML, a two-component phenomenological model was preferred (ΔAICc = −12.1, −24.1 and −4.6, respectively), resolving the response into a higher-sensitivity component carrying half to three quarters of the effect and a lower-sensitivity component five to six orders of magnitude weaker (Supplementary Table 1). The two models were indistinguishable for AMI (ΔAICc = +0.2), and the single-component fit was clearly preferred for ABZ (ΔAICc = +11.6). Two caveats bound these fits. The higher-sensitivity IC_50_ fell at or near the lowest concentration tested, making it an upper bound rather than an estimate, and the confidence intervals of several parameters are wide (Supplementary Table 1); the separation between components should therefore be read as evidence of a biphasic concentration–response profile rather than as a quantitative estimate of two potencies. The intermediate plateau, in turn, sits near 50 % of control, so the IC_50_ shifts with the model chosen, and Table 1 reports 4PL values throughout to keep potencies comparable among compounds.

Across the panel, motility was nearly abolished at concentrations at which most organisms still excluded methylene blue, and lethal concentrations exceeded those required to inhibit motility by more than two orders of magnitude in every compound. CD and AML had the lowest lethal concentrations of the panel and AMI the highest (Table 1).

The compounds differed in efficacy as much as in potency. Ion channel agents removed essentially the entire population at the top of their ranges, whereas the two cytoskeletal modulators plateaued short of complete killing (Table 1). Because the ABZ curve never crossed 50 % lethality within the tested range, its LC_50_ is obtained by extrapolation and its upper confidence limit is not constrained by the data.

### 3.2. Divergent relationships between motility inhibition and loss of viability

To determine whether drug-induced impairment of motility reflected loss of parasite viability, the two endpoints were compared across the full concentration range tested for each compound, taking as the unit of analysis the mean of each compound–concentration combination (n = 39; Table 2, Fig. S1). PZQ, AML and ABZ each showed a significant inverse association between motility inhibition and viability, whereas CD and AMI did not. Coupling between the two endpoints was therefore not a uniform property of either drug class.

**Table 2.** Correlation between motility inhibition (%) and viability (%) after 24 h exposure.

| Treatment | $r_s$ | 95% CI | $p$ value |
| --- | --- | --- | --- |
| Albendazole | -0.89 | [-1.00, -0.40] | 0.037 |
| Cytochalasin D | -0.55 | [-1.00, 0.34] | 0.342 |
| Praziquantel | -0.98 | [-1.00, -0.79] | 0.002 |
| Amlodipine | -0.88 | [-1.00, -0.32] | 0.029 |
| Amiloride | 0.29 | [-0.59, 1.00] | 0.501 |
| Partial correlations | dose | dose + compound |  |
| All compounds | -0.29 ns<br>[-0.62, 0.16] | 0.07 ns<br>[-0.37, 0.49] |  |
| Cytoskeletal modulators | 0.42 ns<br>[-0.71, 0.84] | 0.42 ns<br>[-0.74, 0.84] |  |
| Ion channel agents | -0.38 ns<br>[-0.73, 0.11] | -0.14 ns<br>[-0.64, 0.44] |  |

The correlation coefficient (Spearman's rho, $r_s$ ) usually presents a weak ( $r < 0.5$ ), moderate ( $0.5 < r < 0.8$ ) and strong ( $0.8 < r$ ) relationship. $p$ values for the five per-compound correlations were corrected by the Holm–Bonferroni procedure. ns: not significant. Square brackets denote 95% confidence intervals for $r_s$ (both per-compound and partial correlations).

Both endpoints increased with concentration, so these per-compound correlations largely reflect that shared dependence rather than a link between them. Adjusting for log₁₀[dose] left the association non-significant (rₛ = −0.29; 95 % CI: 0.62 to 0.16), and adjusting additionally for compound identity brought the coefficient close to zero (rₛ = 0.07; 95 % CI: 0.37 to 0.49), with the same pattern within each drug class (Table 2).

The relationship between the two endpoints was not described by a single linear trend (Supplementary Figure 1); viability changed only minimally over an initial range of motility loss and then declined more steeply, a threshold-like pattern captured by segmented regression (Figure 3). In ion channel agents the estimated transition lay at 75.1 % motility inhibition (95 % CI: 71.5– 82.4; p < 0.001), with slopes of −0.06 and −3.88 % viability per unit of motility inhibition below and above it. In cytoskeletal modulators the change in the relationship occurred earlier, at 54.0 % inhibition (95 % CI: 46.6–91.8; p = 0.019), and the subsequent decline was more gradual (−0.14 and −1.02). Individually, AMI, AML and PZQ were biphasic, whereas ABZ and CD were not, which is why the pattern is reported at class level; per-compound estimates are given in Supplementary Table 2.

**Figure 3.**
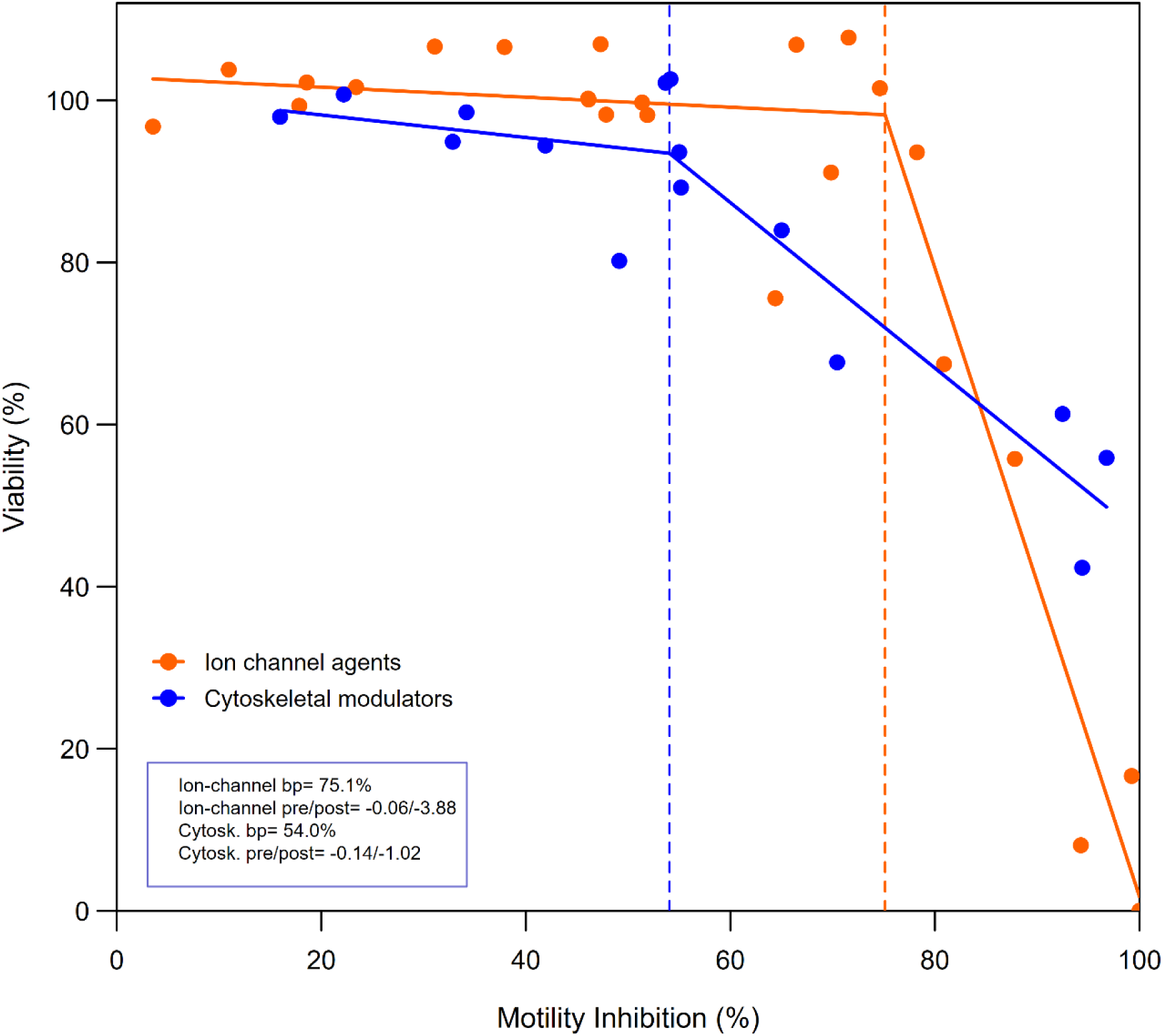
Segmented regression of viability on motility inhibition in *E. granulosus* PSCs. Separate models were fitted to the pooled observations of each drug class: ion-channel agents (orange; n = 24) and cytoskeletal modulators (blue; n = 15). Each point is the mean of one compound–concentration combination. Dashed vertical lines mark the estimated breakpoint of each class; the inset gives breakpoints and the slopes before and after them (viability % per unit of motility inhibition %). Both segmented fits improved significantly on a linear model (p < 0.001 and p = 0.019, respectively).

### 3.3. Persistence of motility impairment after compound removal

To distinguish transient paralysis from sustained impairment, PSCs were washed after the 24 h exposure and maintained for a further 24 h in drug-free medium (Figure 4). Five compound– concentration combinations were excluded because mean post-washout viability fell below 15 %, a level at which the motility readout is no longer reliable; these are reported as not interpretable rather than as evidence for or against recovery.

**Figure 4.**
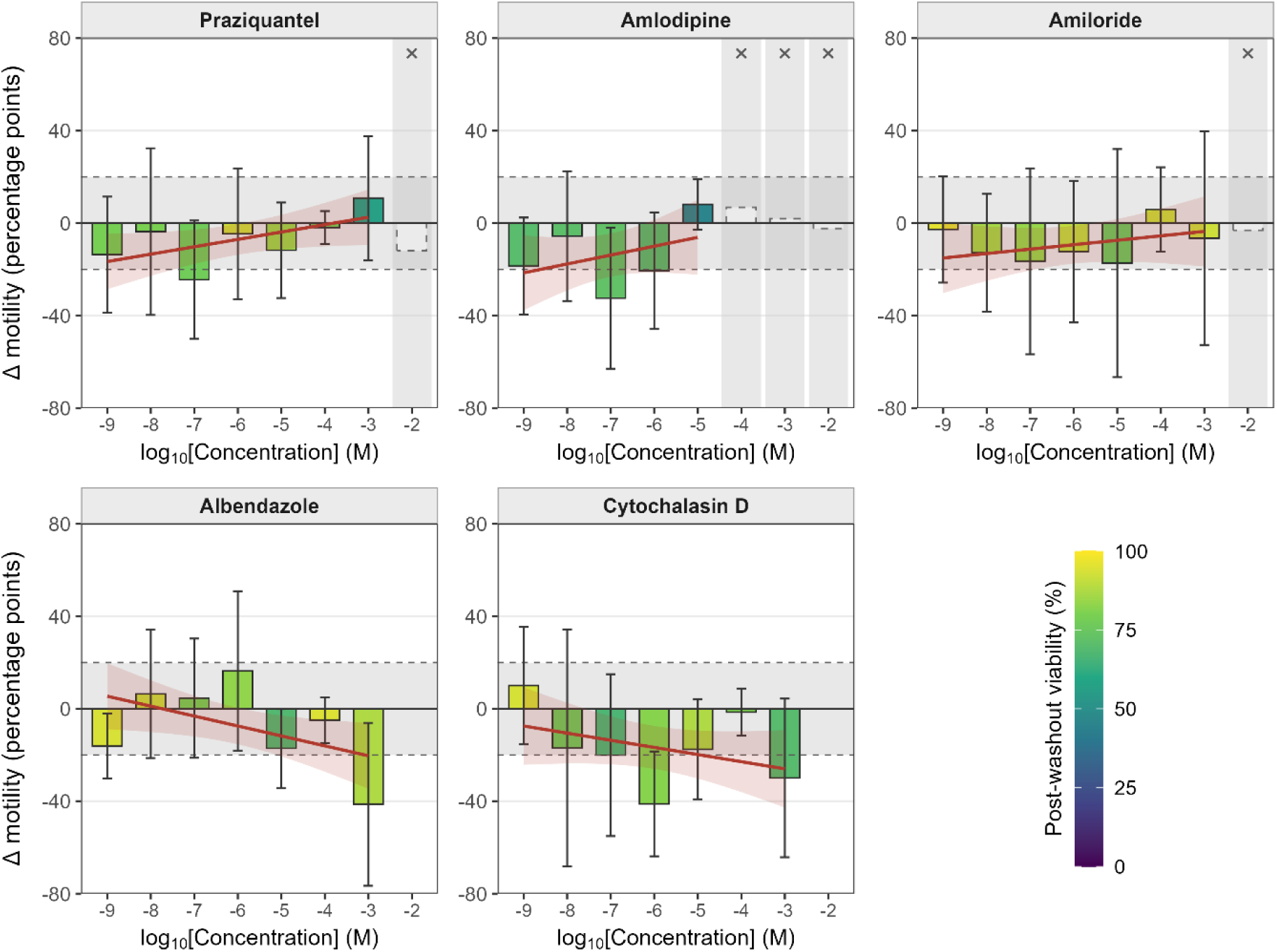
Reversibility of contractile activity in *E. granulosus* PSCs after compound washout. Change in motility (Δ) was calculated well by well as the difference between motility measured 24 h after washout and motility measured after 24 h of treatment and is expressed in percentage points of motility. Bars show the mean Δ for each concentration and error bars show the 95% confidence interval. Δ > 0 indicates that motility increased after washout (recovery), whereas Δ < 0 indicates that motility continued to decline. Red line shows the fitted concentration–recovery trend for each compound with its 95% confidence band. The grey horizontal band delimits the technical-noise band (± 20 percentage points), the standard deviation of the difference between technical replicates in pre-washout values (SD = 19.4 percentage points, n = 109 pairs. Bar fill encodes mean post-washout viability. Shaded columns marked with a cross denote combinations excluded from the analysis because mean post-washout viability fell below 15%.

No compound showed consistent or statistically supported recovery of motility after washout. Across the evaluable combinations Δ scattered around zero or below it, indicating that parasites neither regained movement nor, in most conditions, deteriorated further (Figure 4). Only ABZ showed a significant dose dependence of Δ, and its sign pointed away from recovery: the higher the concentration, the further motility declined after drug removal. The remaining four compounds showed no significant trend (Supplementary Table 3). At 10⁻⁴ M, PZQ, ABZ and CD each produced a Δ near zero while post-washout viability remained between 78 % and 95 % (Figure 4, bar fill), organisms that were alive but immobile.

### 3.4. Ultrastructural damage of PSCs induced by cytoskeletal and ion channel–targeting drugs

Scanning electron microscopy (SEM) showed that all five compounds altered the tegument and the gross morphology of the PSCs, with severity and pattern differing by mechanism of action (Figure 5). Under control conditions, PSCs retained a well-preserved architecture, with clearly defined rostellum, suckers and body, and a continuous tegument densely covered by uniformly distributed microtrichia (Figure 5A). ABZ produced moderate tegumental damage, with partial distortion of the body and suckers and an irregular surface; at higher magnification the tegument was disorganized and had lost microtrichia over part of its extent (Figure 5B). CD, which disrupts actin filaments, produced a more severe phenotype: the suckers and body were deformed and collapsed, the tegument was extensively disrupted, and microtrichia were lost, shortened or disorganized across a surface that frequently appeared corrugated and eroded (Figure 5C).

**Figure 5.**
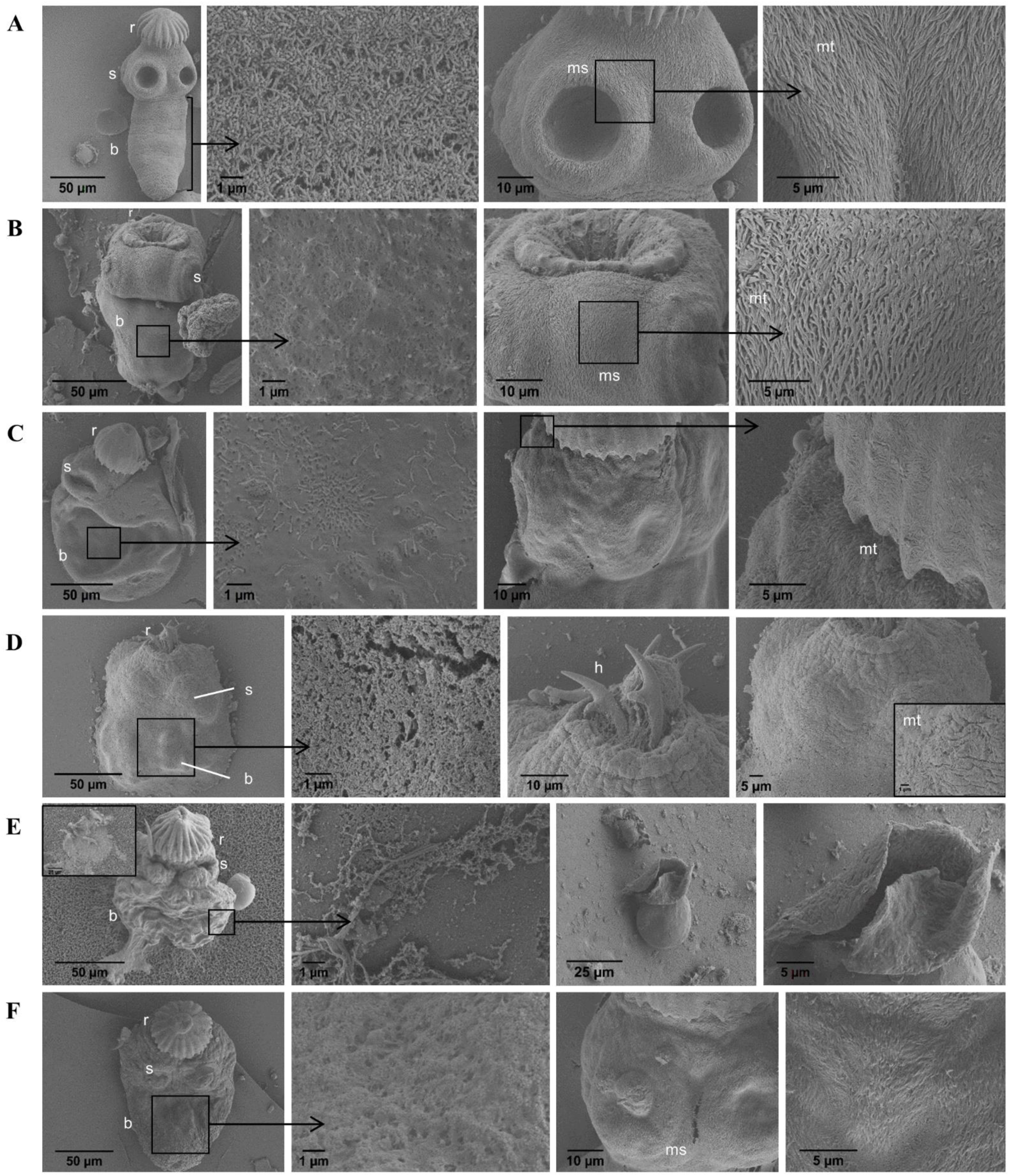
Scanning electron microscopy of *E. granulosus* PSCs treated with cytoskeletal and ion channel drugs. Parasites were incubated with: (A) 0.1 % DMSO (solvent control), (B) 0.1 mM Albendazole (ABZ), (C) 0.1 mM Cytochalasin D (CD), (D) 1 mM Praziquantel (PZQ), (E) 1 mM Amlodipine (AML), and (F) 1 mM Amiloride (AMI). For each condition, panels show the whole parasite (left), tegument surface detail at high magnification (second column), the scolex region (third column), and microtrichia detail (right). r: rostellum; s: sucker; b: body; ms: muscular sucker; mt: microtrichia; h: hooks.

Among the ion channel modulators, PZQ caused the most severe damage. The tegument collapsed, often exposing the underlying syncytial layer, the rostellar pad was distorted, and hooks were displaced or embedded within folded tegument (Figure 5D); microtrichia were truncated or absent. AML also produced substantial alterations — strong body contraction, thick tegumental folds and vesicle-like structures, particularly around the suckers, and a highly disorganized surface with abundant detached debris (Figure 5E). Its microtrichia were sparse, shortened and irregularly arranged. AMI produced the mildest lesions of the panel: an irregular surface with shallow erosions and slight collapse in the rostellar region (Figure 5F). The tegument remained partially preserved, microtrichia were less dense and unevenly oriented, and the rostellar hooks retained their structure.

## 4. Discussion

Cystic echinococcosis remains a neglected tropical disease for which chemotherapy is limited and often ineffective, leaving surgery as the only curative option despite the risks of recurrence and postoperative complications (Kern, 2010; Kern et al., 2017; Prousalidis et al., 2012). This therapeutic gap underscores the importance of phenotypic screening as one route to the discovery of compounds that act rapidly on the parasite. To this end, motility-based assays have become common, because they give an objective, quantifiable readout of neuromuscular activity and overall fitness (Buckingham et al., 2014; Ritler et al., 2017). In the present study, tested compounds targeting two systems on which the PSCs depend, cytoskeletal organization and ion transport, both of which support tegumental architecture, intracellular transport, signaling and host–parasite interaction (Martinac, 2014; Ríos-Valencia et al., 2023; Robertson and Martin, 2007).

All five compounds nearly abolished PSCs motility within the concentration ranges tested, but they differed by more than three orders of magnitude in the concentration required to do so, and by a comparable margin in the concentration required to compromise viability. The two endpoints were not equivalent: in every compound movement was lost well before membrane integrity, and the ordering of compounds by lethal concentration did not follow their ordering by inhibitory concentration.

ABZ and CD act on complementary arms of the cytoskeleton. ABZ targets β-tubulin and inhibits microtubule polymerization, disrupting vesicular transport, glucose uptake and tegumental turnover (Guarnaschelli and Koziol, 2025), whereas CD inhibits actin filament polymerization, a process on which tegumental stability and parasite morphology depend (Ferreira de Lima et al., 2021; La-Rocca et al., 2019; Lacey, 1990). Both drugs impaired motility at submicromolar to micromolar concentrations, but their structural consequences diverged. ABZ produced moderate tegumental disorganization with partial loss of microtrichia, while CD caused pronounced deformation and collapse of the suckers and body together with extensive surface erosion (Figure 5B–C), and was also the more lethal of the two. The contrast is consistent with the two targets: loss of microtubules degrades transport and turnover progressively, whereas loss of the actin cortex removes the element that maintains tegumental shape directly.

For some compounds, particularly PZQ and CD, a two-component model described the data better than one. What those components are, this design cannot say. The compounds may act on more than one target, the PSCs population may be heterogeneous in sensitivity, or two aspects of movement may be lost at different concentrations. ABZ is the only compound in the panel with a single well-characterized target.

Among the ion channel agents, PZQ is widely used against trematodes and many cestodes (Urrea-Paris et al., 1999); in Schistosomes it produces rapid depolarization, sustained muscle contraction and extensive tegumental damage, effects now linked to activation of a TRP channel that mediates non-selective Ca²⁺ influx in excitable cells (Park and Marchant, 2020; Chulkov et al., 2021, 2023). Dihydropyridine Ca^2+^ channel blockers such as AML are not classical anthelmintics, but flatworm neuromuscular systems are highly sensitive to Ca²⁺-channel modulation (Jeziorski and Greenberg, 2006; McVeigh et al., 2005), and AML was the most potent compound of the panel. PZQ and ABZ both deformed the rostellar region preferentially; its dependence on cytoskeletal integrity and on Ca²⁺-regulated muscular activity leaves it vulnerable to either perturbation. AML instead produced strong body contraction with thick tegumental folds and vesicle-like structures around the suckers (Figure 5B, D–E).

AMI was distinct both mechanistically and phenotypically. In vertebrate epithelia it is a prototypical blocker of epithelial Na⁺ channels and Na⁺/H⁺ exchangers (Sariban-Sohraby and Benos, 1986), and comparable sensitivities have been reported in platyhelminths: AMI impairs recovery from acid loads across the tegument of *Schistosoma mansoni* (Pax and Bennett, 1990), and in *E. granulosus* it alters tegumental electrical potentials under high-K⁺, low-Ca²⁺ conditions (Carabajal et al., 2023). Here it was the least potent compound in both endpoints, producing the mildest ultrastructural lesions, with rostellar hooks retaining their structural integrity (Figure 5F). AMI was the only compound in which motility inhibition and viability were unrelated. Taken with its known targets, this profile is consistent with a primary action on electrodiffusional cation movement across the tegument, modulating membrane potential and neuromuscular excitability without a comparable effect on structural integrity.

The correlation and segmented analyses bear on how these phenotypes should be read. Motility impairment generally preceded loss of viability, and the marginal correlations between the two endpoints — strong for PZQ, AML and ABZ — did not survive adjustment for concentration. Loss of motility and loss of viability are better read as separate responses to the same compound, the first occurring well below the concentrations that reduce viability, than as two readouts of a single progressive process; their apparent association across the dose range follows from both being driven by concentration. Where the relationship changed, it changed at a different point in each class, and more abruptly in the ion channel group.

This reading depends on what the viability assay measures. Methylene blue exclusion reports the integrity of the tegumental membrane, not metabolic competence: it does not measure ATP content or metabolic flux, and it will not detect early physiological injury in an organism whose surface remains intact (Casado et al., 1986; Ritler et al., 2017). A compound that arrests movement without permeabilizing the tegument will therefore register as non-lethal whether or not the parasite remains metabolically viable, and the dissociation observed for AMI is open to both readings.

Motility did not recover in any compound after 24 h in drug-free medium (post washout), and for ABZ it continued to decline as concentration increased. The convergence of PZQ, ABZ and CD at 10⁻⁴ M, a persistent motor deficit in parasites that remained largely viable, is informative precisely because these compounds act through distinct primary targets and suggests that the neuromuscular apparatus of the PSCs has limited capacity to recover once perturbed, irrespective of the route by which the perturbation is introduced. The design establishes that no recovery occurred within the 24-h window assayed, not that the deficit is permanent.

The ultrastructural findings run parallel to this. All compounds altered the tegument, most severely PZQ and CD, with ABZ and AML producing intermediate phenotypes and AMI the mildest. Alteration or loss of microtrichia was the one feature common to every treatment, and given the role of these structures in nutrient absorption and host–parasite interaction it may represent a shared route to parasite dysfunction. Whether the organisms left immobile but membrane-intact are metabolically compromised cannot be resolved with the present endpoints and will require further experimentation attending to metabolic activity directly.

## 5. Conclusions

This study shows that motility inhibition and loss of viability in *E. granulosus* protoscoleces reflect largely distinct processes: in both drug classes movement was nearly abolished at concentrations well below those required to compromise membrane integrity, the association between the two endpoints did not survive adjustment for concentration, and the impairment did not reverse within 24 h of washout. Beyond this dissociation, the compound panel points to two convergent routes of vulnerability. Agents that disrupt Ca²⁺-dependent signaling (PZQ and AML) and the actin cytoskeleton (CD) produced the most potent motility inhibition and the most severe tegumental damage. Calcium homeostasis and cytoskeletal integrity thus emerge as the principal physiological pressure points in the protoscolex, and their simultaneous disruption is a rational strategy to test for improved scolicidal efficacy.

## Data Availability Statement

The raw data supporting the conclusions of this article will be made available by the authors, without undue reservation.

## Ethics Statement

Protoscoleces of *E. granulosus* s.l. were obtained from hydatid cysts present in the lungs of naturally infected cattle, collected at a commercial abattoir as a by-product of routine slaughter for human consumption. No live animals were used or manipulated for the purposes of this study and no procedure was performed on animals. Ethical approval was not required for the study involving animals in accordance with the local legislation and institutional requirements because the material analyzed was obtained post-mortem from animals slaughtered for commercial purposes unrelated to this research.

## Author Contributions

HFC had the original project idea and supervised the research. MPAC, MJFS and DDL designed and performed the experiments. MPAC carried out data curation and formal analysis. LJM processed the samples and acquired the bioimages. VHA and EM analyzed the bioimages. HFC and VHA provided essential reagents and equipment. MPAC, HFC and VHA wrote the manuscript, and all remaining authors critically revised it. All authors contributed to the article and approved the submitted version.

## Acknowledgments

We thank Hernán Esquivel and Roberto Fanjul for technical assistance with imaging samples. All micrographs were taken at the Centro Integral de Microscopía Electrónica (CIME), belonging to UNT and CONICET, in Tucumán, Argentina.

A previous version of this manuscript has been released as a preprint at bioRxiv: https://doi.org/10.64898/2026.04.06.716494

## References

1. Alvarez Rojas, C.A., Romig, T., Lightowlers, M.W., 2014. *Echinococcus granulosus* sensu lato genotypes infecting humans–review of current knowledge. Int. J. Parasitol. 44, 9–18. 10.1016/j.ijpara.2013.08.008.

2. Buckingham, S.D., Partridge, F.A., Sattelle, D.B., 2014. Automated, high-throughput, motility analysis in *Caenorhabditis elegans* and parasitic nematodes: Applications in the search for new anthelmintics. Int. J. Parasitol. Drugs Drug Resist. 4, 226–232. 10.1016/j.ijpddr.2014.10.004.

3. Carabajal, M.P.A., Durán, M.A., Olivera, S., Fernández Salom, M.J., Cantiello, H.F., 2022. Electrical potentials of protoscoleces of the cestode *Echinococcus granulosus* from bovine origin. Exp. Parasitol. 238, 108282. 10.1016/j.exppara.2022.108282.

4. Carabajal, M.P.A., Fernández Salom, M.J., Olivera, S., Cantiello, H.F., 2023. Effect of temperature and ionic substitutions on the tegumental potentials of protoscoleces of *Echinococcus granulosus*. Trop. Med. Infect. Dis. 8, 303. 10.3390/tropicalmed8060303.

5. Casado, N., Rodriguez-Caabeiro, F., Hernandez, S., 1986. In vitro survival of *Echinococcus granulosus* protoscoleces in several media, at +4°C and +37°C. Z. Parasitenkd. 72, 273–278. 10.1007/bf00931155.

6. Chen, X.Z., Li, Q., Wu, Y., Liang, G., Lara, C.J., Cantiello, H.F., 2008. Submembranous microtubule cytoskeleton: interaction of TRPP2 with the cell cytoskeleton. FEBS J., 275, 4675– 4683. 10.1111/j.1742-4658.2008.06616.x.

7. Choudhary, S., Kashyap, S.S., Martin, R.J., Robertson, A.P., 2022. Advances in our understanding of nematode ion channels as potential anthelmintic targets. Int. J. Parasitol. Drugs Drug Resist. 18, 52–86. 10.1016/j.ijpddr.2021.12.001.

8. Chulkov, E.G., Smith, E., Rohr, C.M., Yahya, N.A., Park, S.-K., Scampavia, L., Spicer, T.P., Marchant, J.S., 2021. Identification of novel modulators of a schistosome transient receptor potential channel targeted by praziquantel. PLoS Negl. Trop. Dis. 15, e0009898. 10.1371/journal.pntd.0009898.

9. Chulkov, E.G., Palygin, O., Yahya, N.A., Park, S.-K., Marchant, J.S., 2023. Electrophysiological characterization of a schistosome transient receptor potential channel activated by praziquantel. Int. J. Parasitol. 53, 415–425. 10.1016/j.ijpara.2022.11.005.

10. Deplazes, P., Rinaldi, L., Rojas, C.A., Torgerson, P., Harandi, M., Romig, T., Antolova, D., Schurer, J., Lahmar, S., Cringoli, G., 2017. Global distribution of alveolar and cystic echinococcosis. Adv. Parasitol. 95, 315–493. 10.1016/bs.apar.2016.11.001.

11. Doenhoff, M.J., Cioli, D., Utzinger, J., 2008. Praziquantel: mechanisms of action, resistance, new derivatives for schistosomiasis. Curr. Opin. Infect. Dis. 21, 659–667. 10.1097/qco.0b013e328318978f.

12. Ferreira de Lima, N., de Andrade Picanço, G., Ríos Valencia, D.G., López Villegas, E.O., Espinoza Mellado, M.D.R., Ambrosio, J.R., Vinaud, M.C., 2021. Alterations in *Taenia crassiceps* cysticerci cytoskeleton induced by nitazoxanide and flubendazole. Acta Trop. 221, 106027. 10.1016/j.actatropica.2021.106027.

13. Guarnaschelli, I., Koziol, U., 2025. Albendazole specifically disrupts microtubules and protein turnover in the tegument of the cestode *Mesocestoides corti*. PLoS Pathog. 21, e1013221. 10.1371/journal.ppat.1013221.

14. Jeziorski, M.C., Greenberg, R.M., 2006. Voltage-gated calcium channel subunits from platyhelminths: potential role in praziquantel action. Int. J. Parasitol. 36, 625–632. 10.1016/j.ijpara.2006.02.002.

15. Kern, P., 2010. Clinical features and treatment of alveolar echinococcosis. Curr. Opin. Infect. Dis. 23, 505–512. 10.1097/qco.0b013e32833d7516.

16. Kern, P., Da Silva, A.M., Akhan, O., Müllhaupt, B., Vizcaychipi, K., Budke, C., Vuitton, D., 2017. The echinococcoses: diagnosis, clinical management and burden of disease. Adv. Parasitol. 96, 259–369. 10.1016/bs.apar.2016.09.006.

17. Krämer, F., Baneth, G., Dantas-Torres, F., Hamer, S., Lappin, M.R., Otranto, D., Roura, X., Sager, H., Schunack, B., Scorza, V., Traub, R., Geary, T.G., 2025. Resistance of companion animal parasites to antiparasitic drugs. Adv. Parasitol. 128, 35–157. 10.1016/bs.apar.2025.07.002.

18. La-Rocca, S., Farias, J., Chalar, C., Kun, A.E., Fernandez, V., 2019. *Echinococcus granulosus*: Insights into the protoscolex F-actin cytoskeleton. Acta Trop. 199, 105122. 10.1016/j.actatropica.2019.105122.

19. Lacey, E., 1990. Mode of action of benzimidazoles. Parasitol. Today. 6, 112–115. 10.1016/0169-4758(90)90227-u.

20. Martinac, B., 2014. The ion channel-cytoskeleton connection as a potential mechanism of mechanosensitivity. Biochim. Biophys. Acta Biomembr. 1838, 682–691. 10.1016/j.bbamem.2013.07.015.

21. McVeigh, P., Kimber, M.J., Novozhilova, E., Day, T.A., 2005. Neuropeptide signalling systems in flatworms. Parasitol. 131, S41–S55. 10.1017/s0031182005008851.

22. Morseth, D.J., 1967. Fine structure of the hydatid cyst and protoscolex of *Echinococcus granulosus*. J. Parasitol. 53, 312–325. 10.2307/3276582.

23. Nogueira, R.A., Lira, M.G.S., Licá, I.C.L., Frazão, G.C.C.G., dos Santos, V.A.F., Filho, A.C.C.M., Rodrigues, J.G.M., Miranda, G.S., Carvalho, R.C., Nascimento, F.R.F., 2022. Praziquantel: An update on the mechanism of its action against schistosomiasis and new therapeutic perspectives. Mol. Biochem. Parasitol. 252, 111531. 10.1016/j.molbiopara.2022.111531.

24. Park, S.K., Marchant, J.S., 2020. The journey to discovering a flatworm target of Praziquantel: A Long TRP. Trends Parasitol. 36, 182–194. 10.1016/j.pt.2019.11.002.

25. Pax, R.A., Bennett, J.L., 1990. Studies on intrategumental pH and its regulation in adult male *Schistosoma mansoni*. Parasitol. 101, 219–226. 10.1017/s0031182000063265.

26. Perez-Serrano, J., Denegri, G., Casado, N., Bodega, G., Rodriguez-Caabeiro, F., 1995. Anti-tubulin immunohistochemistry study of *Echinococcus granulosus* protoscoleces incubated with albendazole and albendazole sulphoxide in vitro. Parasitol. Res. 81, 438–440. 10.1007/bf00931507.

27. Prousalidis, J., Kosmidis, C., Anthimidis, G., Kapoutzis, K., Karamanlis, E., Fachantidis, E., 2012. Postoperative recurrence of cystic hydatidosis. Can. J. Surg. 55, 15–20. 10.1503/cjs.013010.

28. R Core Team, 2024. R: A Language and Environment for Statistical Computing. R Foundation for Statistical Computing, Vienna, Austria. https://www.R-project.org/

29. Rasuk, M.C., Ferrer, G.M., Kurth, D., Portero, L.R., Farías, M.E., Albarracín, V.H., 2017. UV-Resistant Actinobacteria from High-Altitude Andean Lakes: Isolation, Characterization, Antagonistic Activities. Photochem. Photobiol. 93, 865–880. 10.1111/php.12759.

30. Ríos-Valencia, D.G., Ambrosio, J., Tirado-Mendoza, R., Carrero, J.C., Laclette, J.P., 2023. What about the cytoskeletal and related proteins of tapeworms in the host’s immune response? An Integrative Overview. Pathogens. 12, 840. 10.3390/pathogens12060840.

31. Ritler, D., Rufener, R., Sager, H., Bouvier, J., Hemphill, A., Lundström-Stadelmann, B., 2017. Development of a movement-based in vitro screening assay for the identification of new anti-cestodal compounds. PLoS Negl. Trop. Dis. 11, e0005618. 10.1371/journal.pntd.0005618.

32. Robertson, A.P., Martin, R.J., 2007. Ion-channels on parasite muscle: pharmacology and physiology. Invertebr. Neurosci. 7, 209–217. 10.1007/s10158-007-0059-x.

33. Romig, T., Deplazes, P., Jenkins, D., Giraudoux, P., Massolo, A., Craig, P.S., Wassermann, M., Takahashi, K., De La Rue, M., 2017. Ecology and life cycle patterns of Echinococcus species. Adv. Parasitol. 95, 213–314. 10.1016/bs.apar.2016.11.002.

34. Sariban-Sohraby, S., Benos, D., 1986. The amiloride-sensitive sodium channel. Am. J. Physiol. Cell Physiol. 250, C175–C190. 10.1152/ajpcell.1986.250.2.c175.

35. Sasaki, J.-i., Ramesh, R., Chada, S., Gomyo, Y., Roth, J.A., Mukhopadhyay, T., 2002. The anthelmintic drug mebendazole induces mitotic arrest and apoptosis in non-small cell lung cancer cells by depolymerizing tubulin. Mol. Cancer Ther. 1, 1201–1209.

36. Sielaff, T.D., Taylor, B., Langer, B., 2001. Recurrence of hydatid disease. World J. Surg. 25, 83–86. 10.1007/s002680020011.

37. Urrea-Paris, M., Moreno, M., Casado, N., Rodriguez-Caabeiro, F., 1999. *Echinococcus granulosus*: praziquantel treatment against the metacestode stage. Parasitol. Res. 85, 999–1006. 10.1007/s004360050672.

38. WHO, 2023. Echinococcosis. World Health Organization. https://www.who.int/news-room/fact-sheets/detail/echinococcosis (accessed 15 August 2026).

39. Wolstenholme, A.J., 2011. Ion channels and receptors as targets for controlling parasitic nematodes. Int. J. Parasitol. Drugs Drug Resist. 1, 2–13. 10.1016/j.ijpddr.2011.09.003.

40. Zheng, H., Zhang, W., Zhang, L., Zhang, Z., Li, J., Lu, G., Zhu, Y., Wang, Y., Huang, Y., Liu, J., Kang, H., Chen, J., Wang, L., Chen, A., Yu, S., Gao, Z., Jin, L., Gu, W., Wang, Z., Zhao, L., Shi, B., Wen, H., Lin, R., Jones, M.K., Brejova, B., Vinar, T., Zhao, G., McManus, D.P., Chen, Z., Zhou, Y., Wang, S., 2013. The genome of the hydatid tapeworm *Echinococcus granulosus*. Nat. Genet. 45, 1168–1175. 10.1038/ng.2757.

