## Supplemental Data 1 for "Morphological and Functional Effects of Cytoskeletal and Ion-Channel Agents on the Protoscolex of *Echinococcus granulosus sensu lato*"

### 1 Supplementary Figure

**Supplementary Figure 1. Correlation analysis between motility inhibition (%) and viability (%) for each compound in *E. granulosus* PSCs after 24 h of exposure.** Each point is the mean of one compound–concentration combination; horizontal and vertical bars are the standard errors of the motility inhibition and viability means, respectively. Symbol size increases with concentration, rescaled within each compound so that the smallest symbol denotes that compound's lowest concentration and the largest its highest; color indicates drug class (orange, ion-channel agents; blue, cytoskeletal modulators). The dotted diagonal marks one-to-one coupling (viability equal to residual motility), and the dashed line marks the vehicle control (100% viability).

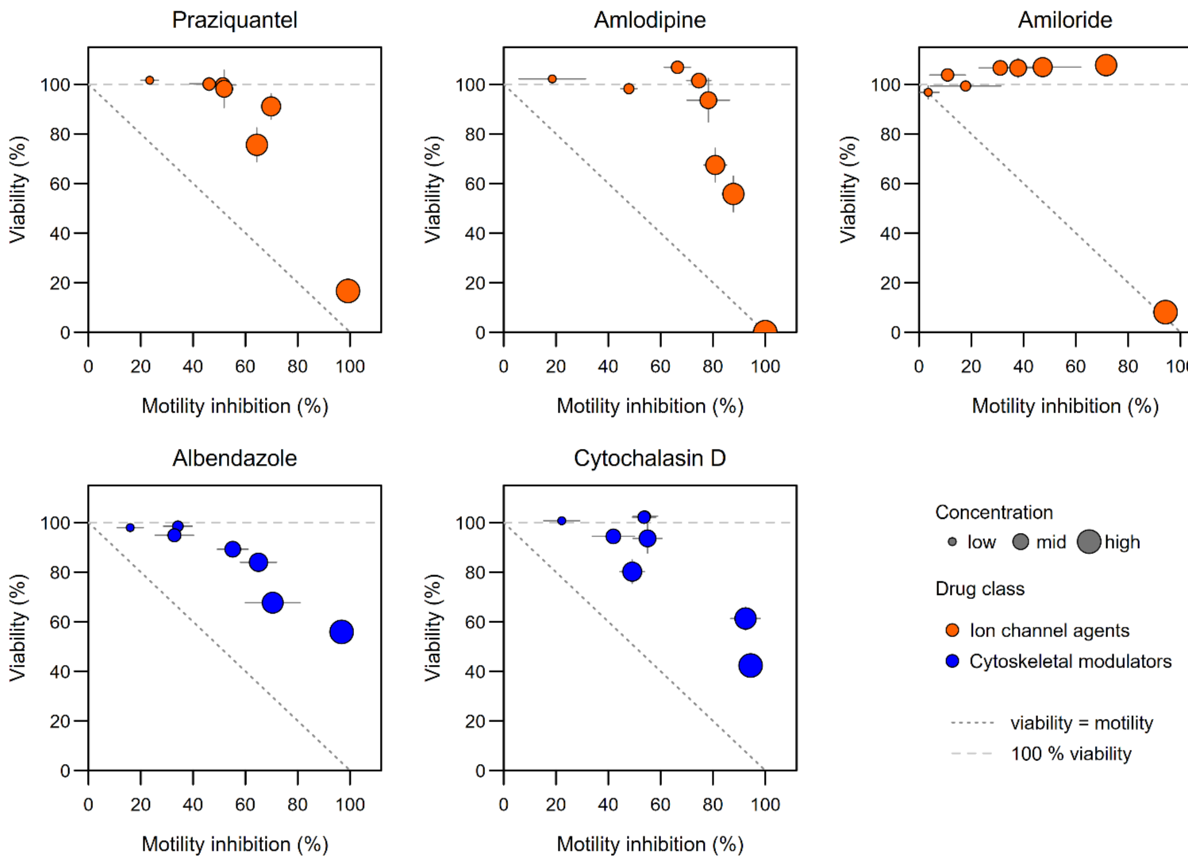

### 2 Supplementary Tables

**Supplementary Table 1.** Comparison of single- and two-component models fitted to the motility concentration–response data.

| Compound | Hill slope [95% CI] | 4PL F (df), p | $IC_{50,1}$, μM [95% CI] | $IC_{50,2}$, μM [95% CI] | F [95% CI] | Two-component F (df), p | ΔAICc (weight) |
|---|---|---|---|---|---|---|---|
| Praziquantel | −0.155 [−0.195, −0.120] | 5.40 (7, 36), < 0.001 | 0.00145 [0.000768–0.00496] | 1810 [721–2920] | 0.54 [0.48–0.60] | 2.92 (6, 36), 0.02 | −12.1 (0.998) |
| Amlodipine | −0.374 [−0.679, −0.200] | 1.97 (5, 31), 0.11 | 0.00493 [0.000657–0.00918] | 638 [1.21–1550] | 0.75 [0.62–0.81] | 0.73 (4, 31), 0.58 | −4.6 (0.909) |
| Amiloride | −0.301 [−0.412, −0.221] | 0.94 (6, 37), 0.48 | 0.209 [0.0014–2.38] | 841 [146–2280] | 0.39 [0.21–0.61] | 0.71 (5, 37), 0.62 | +0.2 (0.478) |
| Albendazole | −0.214 [−0.282, −0.160] | 2.29 (6, 30), 0.06 | $4.40 \times 10^{-5}$ [$1.00 \times 10^{-9}$–0.0146] | 15.9 [1.71–330] | 0.35 [0.25–0.59] | 5.11 (5, 30), 0.002 | +11.6 (0.003) |
| Cytochalasin D | −0.154 [−0.207, −0.108] | 7.19 (7, 37), < 0.001 | $9.93 \times 10^{-5}$ [$2.75 \times 10^{-5}$–0.000205] | 40.8 [23.4–78.2] | 0.50 [0.44–0.56] | 2.02 (6, 37), 0.09 | −24.1 (1.000) |

F is the fraction of the response due to the higher-affinity component. Lack of fit was tested against pure error estimated from replicate wells; a significant F indicates that the model departs from the data by more than experimental variability accounts for. ΔAICc is the two-component minus the four-parameter logistic value, so negative values favor the two-component model; the Akaike weight is its relative probability.

**Supplementary Table 2.** Segmented regression of viability (%) on motility inhibition (%). Each observation is the mean of one compound–concentration combination.

| | Breakpoint (% inhibition) | 95% CI | Flat-profile range | Slope before | Slope after | $R^2$ | *p* value |
|---|---|---|---|---|---|---|---|
| **Cytoskeletal Modulators** | | | | | | | |
| Albendazole | 44.6[b] | [34.2, 70.4] | [40.3, 49.9] | -0.06 | -0.81 | 0.944 | 0.180 |
| Cytochalasin D | 89.4[b] | [49.1, 91.8] | [87.4, 90.6] | -0.10 | -9.90[a] | 0.896 | 0.089 |
| **Ion Channel Agents** | | | | | | | |
| Praziquantel | 69.8 | [46.1, 69.8] | [69.1, 69.8] | -0.42 | -2.40[a] | 0.952 | 0.035 |
| Amlodipine | 74.0 | [69.4, 80.9] | [73.0, 75.1] | 0.07 | -3.96 | 0.979 | 0.004 |
| Amiloride | 70.3 | [10.9, 71.3] | [70.0, 70.7] | 0.22 | -4.39[a] | 0.997 | < 0.001 |
| **Drug-class pooled fits** | | | | | | | |
| All cytoskeletal modulators | 54.0 | [46.6, 91.8] | [52.4, 57.1] | -0.14 | -1.02 | 0.870 | 0.019 |
| All ion channel agents | 75.1 | [71.5, 82.4] | [73.6, 76.9] | -0.06 | -3.88 | 0.938 | < 0.001 |

Flat-profile range: breakpoint values whose residual sum of squares lies within 5% of the minimum, a wide range indicating a breakpoint poorly determined by the data. p values are from a parametric bootstrap under the null hypothesis of linearity. Segmented fits were not corrected for multiple comparisons. [a]Estimated from two observations beyond the breakpoint; the value is unstable and is reported for completeness; it should not be interpreted quantitatively. [b] The segmented model does not fit significantly better than a linear one ($p \geq 0.05$).

**Supplementary Table 3.** Reversibility of contractile activity after compound washout.

| | n wells | n doses | Slope | 95% CI | log[Dose] range | p | Adj. $R^2$ | p (experiment) |
|---|---|---|---|---|---|---|---|---|
| **Cytoskeletal Modulators** | | | | | | | | |
| Albendazole | 42 | 7 | −4.295 | [−8.255, −0.336] | −9 to −3 | 0.034 | 0.215 | 0.015 |
| Cytochalasin D | 42 | 7 | −3.088 | [−7.735, 1.560] | −9 to −3 | 0.187 | 0.116 | 0.048 |
| **Ion Channel Agents** | | | | | | | | |
| Praziquantel | 42 | 7 | 3.192 | [−0.136, 6.520] | −9 to −3 | 0.060 | 0.261 | 0.003 |
| Amlodipine | 30 | 5 | 3.840 | [−2.775, 10.455] | −9 to −5 | 0.243 | 0.051 | 0.227 |
| Amiloride | 40 | 7 | 1.916 | [−2.331, 6.163] | −9 to −3 | 0.366 | 0.303 | < 0.001 |

Δ = motility measured post-washout − motility measured pre-washout, computed well by well so that each well serves as its own control. Data shown are those plotted in Figure 4.
